# Between-area communication through the lens of within-area neuronal dynamics

**DOI:** 10.1101/2022.04.11.487906

**Authors:** Olivia Gozel, Brent Doiron

**Affiliations:** Departments of Neurobiology and Statistics, University of Chicago, Chicago, IL 60637, USA; Grossman Center for Quantitative Biology and Human Behavior, University of Chicago, Chicago, IL 60637, USA

## Abstract

A core problem in systems and circuits neuroscience is deciphering the origin of shared dynamics in neuronal activity: do they emerge through local network interactions, or are they inherited from external sources? We explore this question with large-scale networks of spatially ordered spiking neuron models where a downstream network receives input from an upstream sender network. We show that linear measures of the communication between the sender and receiver networks can discriminate between emergent or inherited population dynamics. Faithful communication requires a match in the dimensionality of the sender and receiver population activities, along with an alignment of their shared fluctuations. However, a nonlinear mapping between the sender – receiver activity or downstream emergent population-wide fluctuations can impair linear communication. Our work exposes the benefits and limitations of linear measures when analyzing between-area communication in circuits with rich population-wide neuronal dynamics.

## Introduction

The brain is composed of a multitude of distributed areas which interact to support the complex computations needed for perception and cognition. While past experimental investigations were typically limited to single neuron recordings, recent technological advances allow for sampling from large populations of neurons simultaneously (Urai et al., 2022). This newfound ability was initially used to characterize the local dynamics of a population of neurons from the same brain area (Schölvinck et al., 2015; Murray et al., 2017; Williamson et al., 2019; Jiang et al., 2020; Langdon et al., 2023). Presently, many studies measure population activity distributed over several brain regions, giving a more holistic, brain-wide view of neuronal processing (Steinmetz et al., 2018; Yu et al., 2019; Musall et al., 2019). However, despite these richer datasets, the science of the mechanics by which different brain areas communicate with one another is still in its infancy.

An often used measure of neuron-to-neuron interaction is the joint trial-to-trial covariability, or noise correlation, of their spike train responses (Cohen and Kohn, 2011; Doiron et al., 2016). The idea is that neuron pairs that have high noise correlations are likely members of the same putative neuronal circuit (Abeles, 1991; Shadlen and Newsome, 1998; Ocker et al., 2017). While pairwise correlations can be informative (Doiron et al., 2016), the large-scale nature of population recordings presents a challenge when attempting to expose the salient aspects of population-wide interactions simply from an analysis of neuron pairs (Das and Fiete, 2020). Dimensionality reduction techniques have been developed to frame population activity within a space of the appropriate size: large enough to capture the core shared variability across a population, yet small enough to be tractable (Cunningham and Yu, 2014; Williamson et al., 2019). These analysis techniques identify low dimensional structure in the population-wide activity, and recent work has used them to measure how connected brain-areas interact with one another (Semedo et al., 2019; Srinath et al., 2021). Yet these techniques do not on their own provide insight into the circuit mechanisms that support or impede brain-area to brain-area communication.

The propagation of brain activity has been the focus of extensive circuit modeling attempts. Feedforward networks are the base structure of many contemporary models of object classification, and have been used with great success to model the performance of visual system hierarchy (Yamins and DiCarlo, 2016). However, networks of spiking neuron models with random, sparse feedforward connectivity produce propagation that leads to excessive, often rhythmic, synchronization (Abeles, 1991; Diesmann et al., 1999; Reyes, 2003; Kumar et al., 2010; Rosenbaum et al., 2011). By contrast, a single population of spiking neuron models with sparse, yet strong, excitatory and inhibitory recurrent connections can show temporally irregular, roughly asynchronous spiking dynamics (Van Vreeswijk and Sompolinsky, 1998; Amit and Brunel, 1997; Renart et al., 2010; Rosenbaum et al., 2017), mimicking what is often considered the default state of cortical networks (Shadlen and Newsome, 1998; Renart et al., 2010). However, neurophysiological recordings over a range of sensory and cognitive states show a wide distribution of spike count correlations whose average is low, but positive and significantly different from zero (Cohen and Kohn, 2011; Doiron et al., 2016). Recent modeling work shows how population dynamics with stable firing rates yet moderate population-wide noise correlations can be produced when structured synaptic wiring is considered, such as discrete block structure (Darshan et al., 2018), low-rank recurrent components (Landau and Sompolinsky, 2018; Mastrogiuseppe and Ostojic, 2018), or distance dependent connection probability (Keane and Gong, 2015; Rosenbaum et al., 2017; Huang et al., 2019). While these results provide new insights into how circuit structure determines shared variability, they have been restricted to within-population dynamics. On the other hand, recent modeling efforts have shed some light on the interaction between brain-areas (Chaudhuri et al., 2015; Muller et al., 2018; Hahn et al., 2019), yet often without a consideration of response fluctuations. Thus, there remains a gap in understanding how the circuit based theories of shared variability within a population extend to the distribution (or propagation) of variability between populations.

In this work, we investigate how complex within brain-area neuronal dynamics affect inter-actions between distinct brain-areas using a network of model spiking neurons with biologically plausible synaptic connectivity and dynamics. We determine conditions when communication is disrupted between an upstream sender network and a downstream receiver network, as assessed by a linear measure. We show that the emergence of complex spatio-temporal dynamics within the sender network leads to faithful sender – receiver communication, while if the receiver generates complex dynamics then communication is disrupted. We understand this dichotomy through how shared fluctuations in the receiver aligns or misaligns with respect to the fluctuations in the sender. Finally, when sender – receiver linear communication is disrupted it occurs in one of two ways: by inducing a nonlinear mapping of the sender – receiver activity, or by yielding chaotic spiking dynamics at the macroscopic scale in the receiver. These results expose the limitations of linear measures when deciphering between brain-area communication in the presence of complex spatio-temporal neuronal dynamics.

## Results

### Destabilization of the E/I balance yields rich within-area population-wide dynamics

Before we explore communication between brain-areas, we first discuss how within brain-area neuronal dynamics depend on the temporal and structural makeup of the recurrent synaptic interactions between neurons. We use a layered network of spiking neuron models that are spatially organized on a square grid (Fig. 1a; see Methods). Neurons in the input layer are modeled as independent homogeneous Poisson processes with a uniform rate. They project their activity to a recurrently coupled network of excitatory (E) and inhibitory (I) spiking neuron models (exponential integrate-and-fire; see Methods). Throughout our study the spatial and temporal scales of synaptic interactions are key parameters. Synaptic currents are modeled as a difference in exponentials with rise and decay timescales, *τ* ^rise^ and *τ* ^decay^, respectively. It is well known that synaptic connectivity is spatially structured with connection probabilities falling off with the distance between pre- and post-synaptic neurons (Holmgren et al., 2003; Levy and Reyes, 2012; Rossi et al., 2020). Accordingly, following past work (Rosenbaum and Doiron, 2014; Rosenbaum et al., 2017; Huang et al., 2019) we model within- and between-layer connectivity as obeying a two-dimensional Gaussian whose spatial widths are denoted by *σ* (see Methods), and we assume periodic boundary conditions on our domain. We set a larger width for the recurrent than the feedforward connections (*σ*_rec_ *> σ*_ffwd_), because such networks exhibit correlated neuronal spiking dynamics (Rosenbaum et al., 2017).

**Figure 1:**
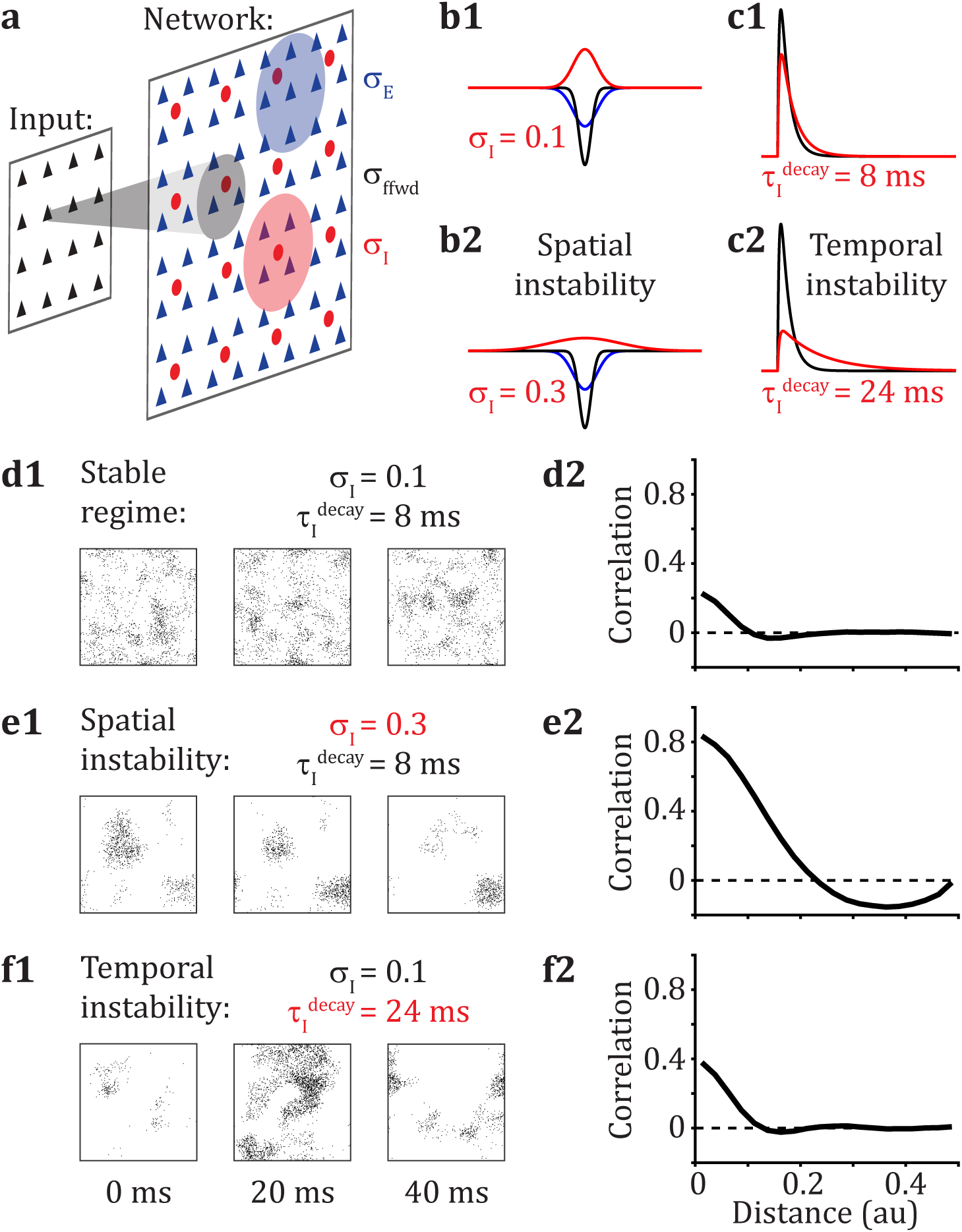
Destabilization of the E/I balance yields rich within-area population-wide dynamics. (**a**) The input layer (black triangles) produces temporally and spatially homogeneous, independent Poisson spike trains. The input layer connects to a network with excitatory (E, blue triangles) and inhibitory (I, red circles) spiking neuron models which are recurrently connected to one another. The neurons are arranged on a two-dimensional [0, 1] *×* [0, 1] grid. Connections are spatially organized according to a wrapped Gaussian (periodic boundary conditions) with widths *σ*_ffwd_, *σ_E_* and *σ_I_* for the feedforward and recurrent E and I connections respectively (standard parameters: *σ*_ffwd_ = 0.05, *σ_E_* = 0.1). (**b**) Connection probability as a function of distance from feedforward input (black), recurrent excitation (blue) and recurrent inhibition (red). In the standard network, recurrent inhibitory connections balance excitatory connections to yield a stable regime (b1, *σ_I_* = 0.1). When recurrent inhibitory connections are broadened (b2, *σ_I_* = 0.3), the E/I balance is spatially destabilized. (**c**) Excitatory post-synaptic potential (EPSP, black) and inhibitory post-synaptic potential (IPSP, red). In the standard network, the timecourse of EPSP and IPSP balance each other to yield a stable regime (c1, *τ_I_* ^decay^ = 8 ms). When the time constant of inhibitory neurons is increased (c2, *τ _I_*^decay^ = 24 ms), the E/I balance is temporally destabilized. (**d1,e1,f1**) Raster plot snapshots of the spiking activity in the recurrent network over 2 ms time-windows separated by 20 ms: (d1) standard network in the stable regime, (e1) spatial instability induced by lateral inhibition, and (f1) temporal instability induced by slow inhibition. (**d2,e2,f2**) Corresponding pairwise correlation as a function of pairwise distance.

Unless otherwise specified, our network is set with the parameters reported in Table 1. We call these the standard parameters (Fig. 1b1,c1) because they yield stable spiking dynamics, as reflected by temporally irregular spiking activity (Fig. 1d1) and an average pairwise spike-count correlation over all spatial scales close to zero (Fig. 1d2) (Rosenbaum et al., 2017). It is known from previous work that when the E/I network parameters lead to destabilization of firing rate dynamics, spatio-temporal patterns of spiking activity intrinsically emerge within the network (Huang et al., 2019; Keane and Gong, 2015; Muller et al., 2018). The E/I balance can either be destabilized in space by increasing the width of recurrent inhibition *σ_I_* (Fig. 1b2,e), or in time by increasing the inhibitory synaptic time constant *τ_I_* ^decay^ (Fig. 1c2,f). The spatio-temporal characteristics of the emerging patterns of activity within the recurrent network depend on the route (spatial or temporal) to E/I destabilization. If E/I balance is spatially destabilized, it yields spatially organized patterns of activity whose spatial scale depends on the width of recurrent inhibition (Fig. 1e1). It has the effect to increase the pairwise spike-count correlations for short pairwise distances, while yielding negative correlations at broader distances (Fig. 1e2). Alternatively, if E/I balance is temporally destabilized, it yields temporally organized patterns of activity that propagate across the entire network (Fig. 1f1). Contrary to a spatial destabilization, the pairwise spike-count correlations are only slightly increased at short pairwise distances (Fig. 1f2). In sum, this modeling framework gives us control of the emergence and structure of complex, population-wide spiking dynamics in the recurrent network without the need for a structured external source of noise. We next investigate if and how complex within-area neuronal dynamics affect the communication between brain-areas.

**Table 1:** Standard parameters for the simulations

|  |  |  |  |  |
| --- | --- | --- | --- | --- |
| Network connectivity | $\sigma_{\text{ffwd}} = 0.05$ | $\sigma_{\text{rec}} = 0.1$ | $\sigma_E = 0.1$ | $\sigma_I = 0.1$ |
| | $\bar{p}_E^{\text{ffwd}} = 0.1$ | $\bar{p}_I^{\text{ffwd}} = 0.05$ | $j_E^{\text{ffwd}} = 240$ | $j_I^{\text{ffwd}} = 400$ |
| | $\bar{p}_{EE}^{\text{rec}} = 0.01$ | $\bar{p}_{EI}^{\text{rec}} = 0.04$ | $\bar{p}_{IE}^{\text{rec}} = 0.03$ | $\bar{p}_{II}^{\text{rec}} = 0.04$ |
| | $j_{EE}^{\text{rec}} = 80$ | $j_{EI}^{\text{rec}} = -240$ | $j_{IE}^{\text{rec}} = 40$ | $j_{II}^{\text{rec}} = -300$ |
| Neuronal dynamics | $C_m = 1 \text{ ms}$ | $g_L^E = 1/15$ | $g_L^I = 1/10$ | $V_L = -60 \text{ mV}$ |
| | $V_{\text{th}} = -10 \text{ mV}$ | $V_{\text{reset}} = -65 \text{ mV}$ | $t_{\text{ref}}^E = 1.5 \text{ ms}$ | $t_{\text{ref}}^I = 0.5 \text{ ms}$ |
| | $V_T = -50 \text{ mV}$ | $\Delta_T^E = 2 \text{ mV}$ | $\Delta_T^I = 0.5 \text{ mV}$ | |
| | $\tau_E^{\text{rise}} = 1 \text{ ms}$ | $\tau_E^{\text{decay}} = 5 \text{ ms}$ | $\tau_I^{\text{rise}} = 1 \text{ ms}$ | $\tau_I^{\text{decay}} = 8 \text{ ms}$ |

### Between brain-area spike count correlations are differentially affected by the location and type of destabilization of the E/I balance

We extend our framework to next explore the responses of a three-layer network of spiking neuron models (Fig. 2a). As before, neurons in the input layer are modeled as independent homogeneous Poisson processes with a uniform rate. The sender (second layer) and receiver (third layer) populations each consist of excitatory (E) and inhibitory (I) spiking neuron models which are spatially organized on a square grid (see Methods). The sender layer projects excitatory connections to the receiver layer, while projections from the receiver to the sender are omitted.

**Figure 2:**
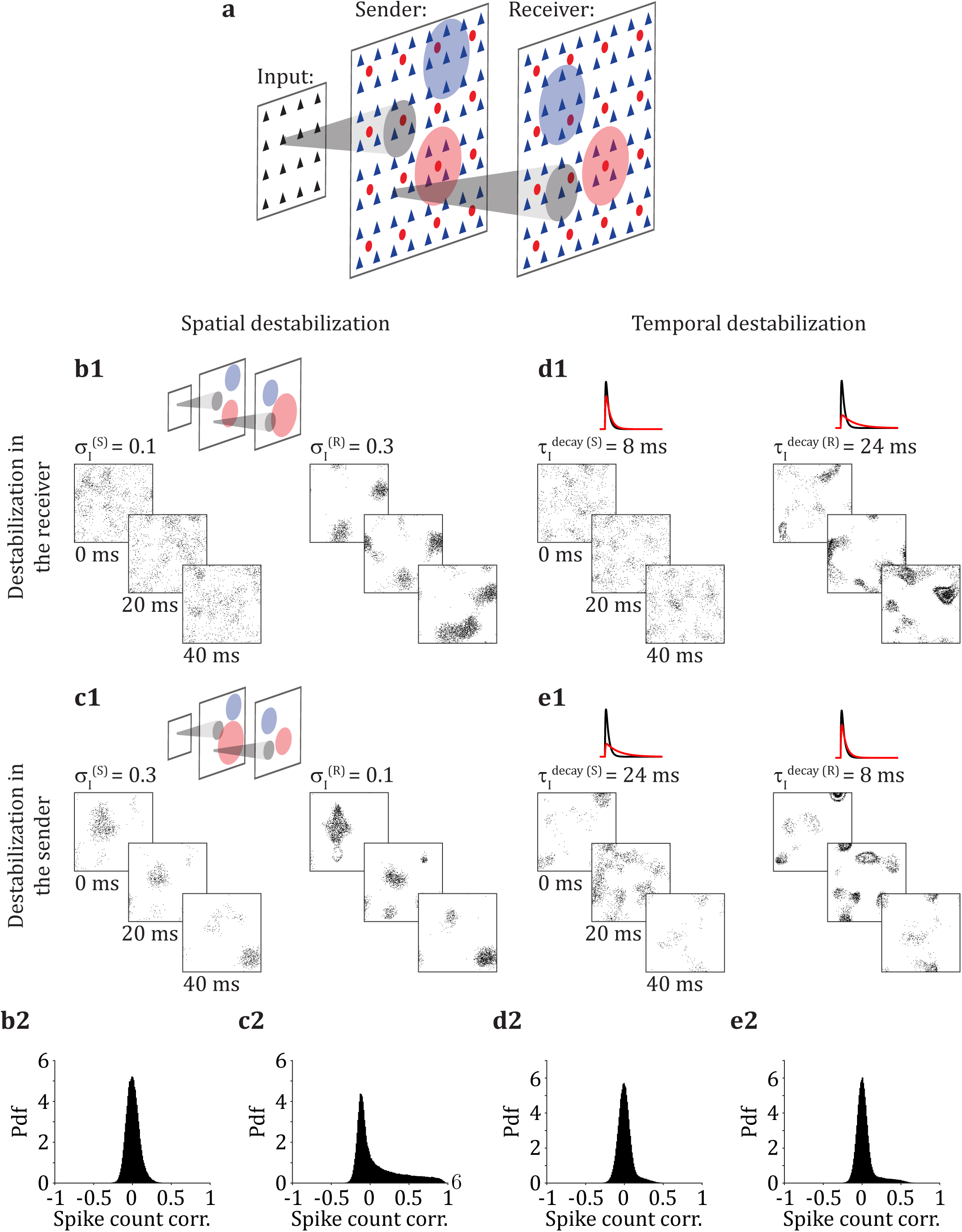
Between brain-area spike count correlations are differentially affected by the location and type of destabilization of the E/I balance. (**a**) The input layer produces homogeneous Poisson spike trains and connects to the sender layer (S), which itself connects to the receiver layer (R). (**b1,c1**) Simultaneous rasters of activity in sender (S, left) and receiver (R, right) layers when the E/I balance is spatially destabilized by increasing the width of recurrent inhibitory connections, *σ_I_*, in R (b1) or in S (c1). (**b2,c2**) Probability density function of the pairwise spike count correlations between a neuron in S and a neuron in R during spatial destabilization in R (b2, mean of the distribution: *µ* = 0.0061) or in S (c2, *µ* = 0.0846). (**d1,e1**) Simultaneous rasters of activity in sender (S, left) and receiver (R, right) layers when the E/I balance is temporally destabilized by increasing the time constant of inhibitory neurons, *τ_I_*^decay^, in R (d1) or in S (e1). (**d2,e2**) Probability density function of the pairwise spike count correlations between a neuron in S and a neuron in R during temporal destabilization in R (d2, *µ* = 0.0052) or in S (e2, *µ* = 0.0304).

The sender network with standard parameters produces spiking activity that is temporally irregular (Fig. 2b1,d1; left), with a near symmetric distribution of pairwise correlations having a mean close to zero (Fig. S1a,c; left). Emergence of spatio-temporal patterns in the receiving network through a spatial destabilization of E/I balance (Fig. 2b1, right) induces a broader distribution of spike count correlations with a heavy positive tail (Fig. S1a, right). Yet the distribution of pairwise correlations between neurons from the sending and receiving networks remains relatively narrow with a mean close to zero (Fig. 2b2). By contrast, when E/I balance is instead spatially destabilized in the sender network, it yields the emergence of spatio-temporal patterns which are subsequently propagated to the receiving network (Fig. 2c1). Consequently the distribution of spike count correlations is heavily-tailed in both the sender and receiver networks (Fig. S1b). Furthermore, we observe a heavy tail in the distribution of between-area spike count correlations (Fig. 2c2).

When E/I balance is destabilized temporally by increasing the time constant of inhibitory synaptic currents, it yields temporally organized patterns of activity. When these patterns originate in the sender network they then propagate downstream to the receiver network (Fig. 2e1). Alternatively, when the receiver network is temporally destabilized then these patterns occur only in the receiver (Fig. 2d1). Despite this patterning, the within-area pairwise spike count correlations increase only slightly (Fig. S1c,d). Further, the between-area spike count correlations also stay low no matter the location (sender or receiver) of the emergence of spatio-temporal patterns (Fig. 2d2,e2). These observations emphasize the diverse characteristics of spatio-temporal population spiking dynamics which depend on the location of their emergence (sender vs. receiver) and the mechanisms through which they are induced (spatial vs. temporal).

### Location and characteristics of spatio-temporal dynamics determine within-area dimensionality

While the average pairwise spike count correlation can be a signature of the dynamical regime of a network (Huang, 2021), we take advantage of our access to large amounts of synthetic data to move beyond statistics on just the bulk pairwise neuronal activity. In this section, we investigate the structure of population-wide shared neuronal variability and how it depends on E/I destabilization.

We randomly select 50 neurons from a local portion of the grid delineated by a disc (Fig. 3a). By changing the radius of the disc we explore how shared variability depends on the spatial scale of the sampled population. We only select neurons whose average firing rate is sufficiently responsive (above 2 Hz) and compute their full spike count covariance matrix *C*. Through Factor Analysis (FA) (Everitt, 1984; Yu et al., 2009), we separate *C* into a shared component, *C*_shared_, and a private component, *C*_private_ (Fig. 3a; see Methods). FA, in contrast to Probabilistic Principal Component Analysis (PPCA), does not assume isotropic noise. Hence the elements of the diagonal matrix *C*_private_ are not constrained to be identical. FA thus determines the directions of highest covariances, and not largest individual variances as in PPCA. Singular Value Decomposition is then applied to *C*_shared_ to obtain the shared eigenvectors and associated eigenvalues which characterize the structure of the shared fluctuations.

**Figure 3:**
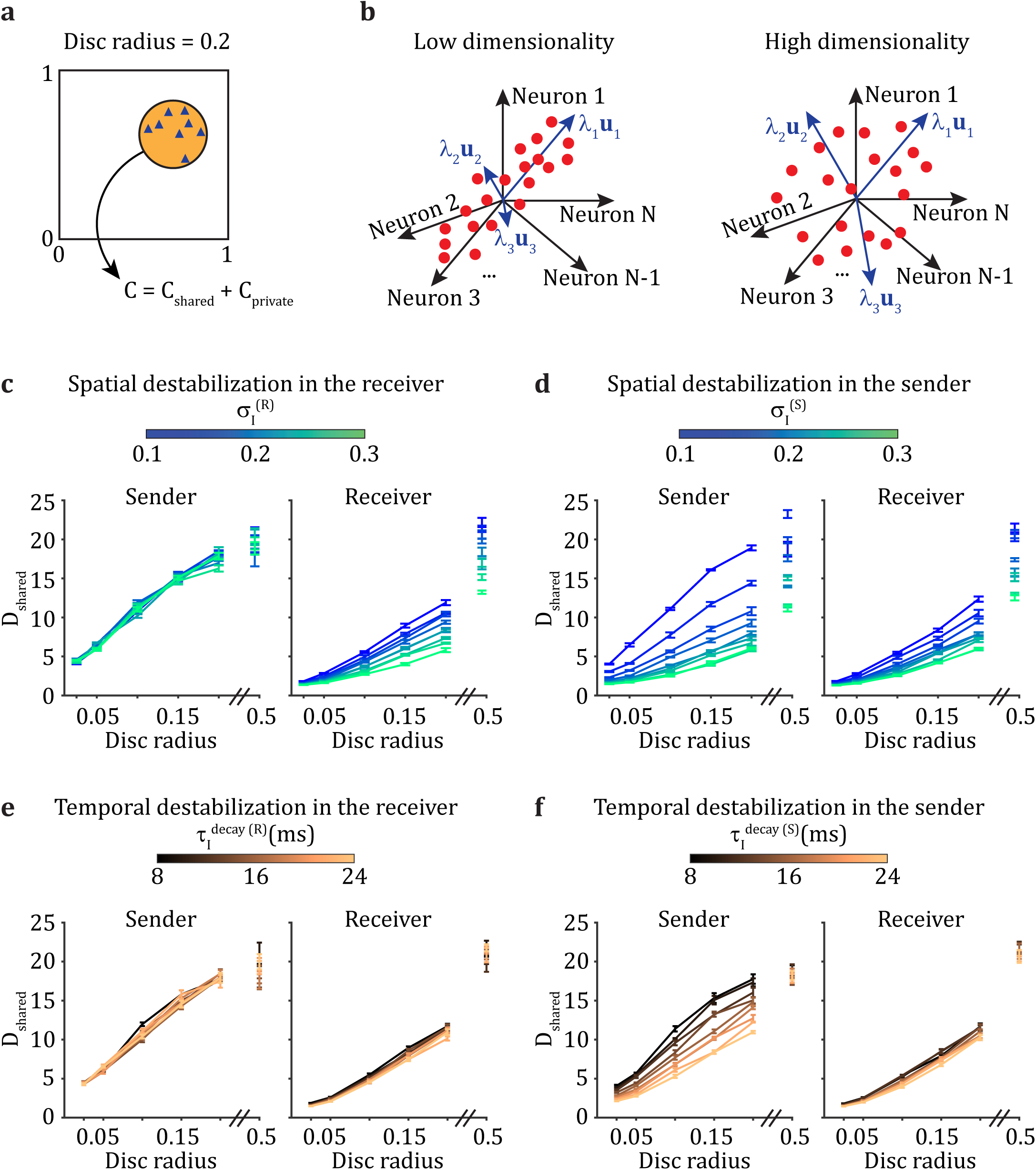
Location and characteristics of spatio-temporal dynamics determine within-area dimensionality. (**a**) Fifty neurons are selected from the neuronal grid by random sampling from discs with different radii. The shared covariance matrix, *C*_shared_, is obtained from the full covariance matrix, *C*, through Factor Analysis. (**b**) Shared dimensionality, *D*_shared_, is measured from the eigenspectrum of *C*_shared_. When the eigenvalues are heterogeneous in magnitude dimensionality is low, whereas when eigenvalues are more uniform in magnitude dimensionality is high. (**c,d**) Shared dimensionality as a function of the radius of the disc from which neurons are sampled, either from the sender (S, left) or from the receiver (R, right), when we destabilize network activity spatially by modifying *σ_I_* in R (c) or in S (d). (**e,f**) Same as (c,d), when we destabilize the network activity temporally by increasing *τ_I_* ^decay^ in R (e) or in S (f).

While the full distribution of the eigenvalues *λ_i_* of the shared covariance matrix *C*_shared_ is informative (Fig. S2, Fig. S3), we want to describe this distribution with a single scalar measure. To this end we compute the *effective dimension* of shared variability from *{λ_i_}* via *D*_shared_ = (Σλ_i_)^2^/Σλ_i_^2^, which is sometimes termed the participation ratio (Mazzucato et al., 2016; Litwin-Kumar et al., 2017). Contrary to other measures of dimensionality (Mante et al., 2013; Kaufman et al., 2014; Williamson et al., 2016; Semedo et al., 2019), *D*_shared_ does not require an arbitrary threshold to give an integer value of dimension. Rather, *D*_shared_ is the squared first moment of the eigenspectrum normalized by the second moment. If the shared fluctuations preferentially take place over a few dimensions, as reflected by a few eigenvalues, *λ_i_*, which are much larger than the others, it yields low dimensionality *D*_shared_ (Fig. 3b, left). On the other hand, if the shared fluctuations are broadly distributed over the whole eigenspace, as reflected by a uniform distribution of the eigenvalues, the resulting dimensionality *D*_shared_ is high (Fig. 3b, right). Using within-area shared dimensionality *D*_shared_, we investigate how the emergence of spatio-temporal patterns affects the structure of shared variability, both within brain-area and in the interaction between connected brain-areas.

In the network with standard parameters, as the disc radius over which neurons are sampled increases the estimated shared dimensionality *D*_shared_ increases (Fig. 3c and e, left). This is expected given how the dominant eigenmode changes with disc radius (Fig. S2a1,b1). However, when sampling neurons from the whole grid (disc radius = 0.5), the shared dimensionality stays much lower than the theoretical upper bound of 50 (Fig. 3c and e, left). Because dimensionality depends strongly on the spatial arrangement of the sampled neurons, we cannot simply associate a unique dimensionality value to the network dynamics as a whole. Instead, we focus on how changes in network dynamics owing to E/I destabilization are reflected in changes in dimensionality.

In the network with standard parameters yielding stable activity in both the sender and the receiver networks, the dimensionality decreases (at fixed disc radius) as activity propagates from the sender to the receiver (*σ_I_*^(R)^ = 0.1 curves in Fig. 3c and *τ _I_*^decay^ ^(R)^ = 8 ms curves in Fig. 3e). When E/I balance is spatially destabilized in the receiver network by increasing the breadth of recurrent inhibitory connections, *σ_I_*^(R)^, spatio-temporal patterns emerge in the receiver yielding a further decrease in dimensionality when compared to that of the sender network (Fig. 3c). By contrast, when E/I balance is instead spatially destabilized in the sender by increasing *σ_I_*^(S)^, then in addition to decreasing within-sender dimensionality locally, the decrease is propagated in the receiver (Fig. 3d).

Different results are obtained if E/I balance is temporally destabilized. When the time constant of the inhibitory synapses, *τ_I_*^decay^, is increased in the receiver network, there is not much change in the dimensionality of the receiver activity (Fig. 3e, right). This suggests that global measures of shared fluctuations cannot detect a temporal destabilization in the receiver, since they yield similar results to the standard network with stable activity. However, if E/I balance is instead temporally destabilized in the sender by increasing *τ_I_* ^decay^ ^(S)^, a decrease in dimensionality is observed in the sender (Fig. 3f, left). Interestingly, this decrease in dimensionality is not propagated to the receiver: all curves overlap, no matter the extent of spatio-temporal patterns inherited from the sender (Fig. 3f, right). The differences in the sensitivity of dimensionality on temporal destabilization between the sender and receiver networks may be due to the independent Poisson input given to the sender from the input layer, while the receiver is given temporally and spatially correlated spiking activity from the sender.

A critical point to remark is that the shared dimensionality in the receiver network is low irrespective if spatio-temporal patterns emerge locally within the receiver network or if they are inherited from the sender network (compare the right panels of Fig. 3c and Fig. 3d, as well as of Fig. 3e and Fig. 3f). This raises an important dilemma for the interpretation of changes in the structure of population-wide shared fluctuations. Namely, that a change in the dimension of population activity can be due to either a shift of the internal dynamics within a network or be inherited from shifts in the dynamics of upstream areas. This ambiguity prompts us to next consider how the sender and receiver networks directly communicate their shared fluctuations.

### Inter brain-area communication strength depends on the origin of shared fluctuations

To measure the interaction between sending and receiving networks we use a recently developed measure of brain-area to brain-area communication based on reduced-rank regression (Semedo et al., 2019). Briefly, the activity in the receiver network, *R*, is predicted from the activity in the sender network, *S*, through a linear model: *R̂* = *SB*_RRR_. The rank of the regression matrix *B*_RRR_ is constrained to be a low value *m* (see Methods). Prediction performance is given by the comparison between *R* and *R̂*, quantifying the ability of the sender population activity in linearly predicting the receiver population activity through a low-dimensional communication subspace.

When E/I balance is destabilized by increasing the breadth of recurrent inhibitory connections in the receiver, given by *σ_I_*^(R)^, the prediction performance of the communication subspace decreases (Fig. 4a). We note that the exact value of prediction performance depends on the spatial scale from which neurons are sampled: the larger the spatial scale, the lower the prediction performance (Fig. S4). By contrast, when E/I balance is instead spatially destabilized in the sender by increasing *σ_I_*^(S)^, the prediction performance of the communication subspace increases (Fig. 4b). If E/I balance is temporally destabilized by increasing the time constant of inhibitory neurons, *τ_I_*^decay^, in either the sender or receiver, qualitatively similar results are obtained (Fig. 4c,d). In sum, despite similar dynamics in the receiver population regardless of the origin of E/I destabilization (Fig. 3c,d,e,f), the prediction performance of the linear communication subspace is unambiguous to the origin: receiver network destabilization disrupts communication, while sender network destabilization improves communication.

**Figure 4:**
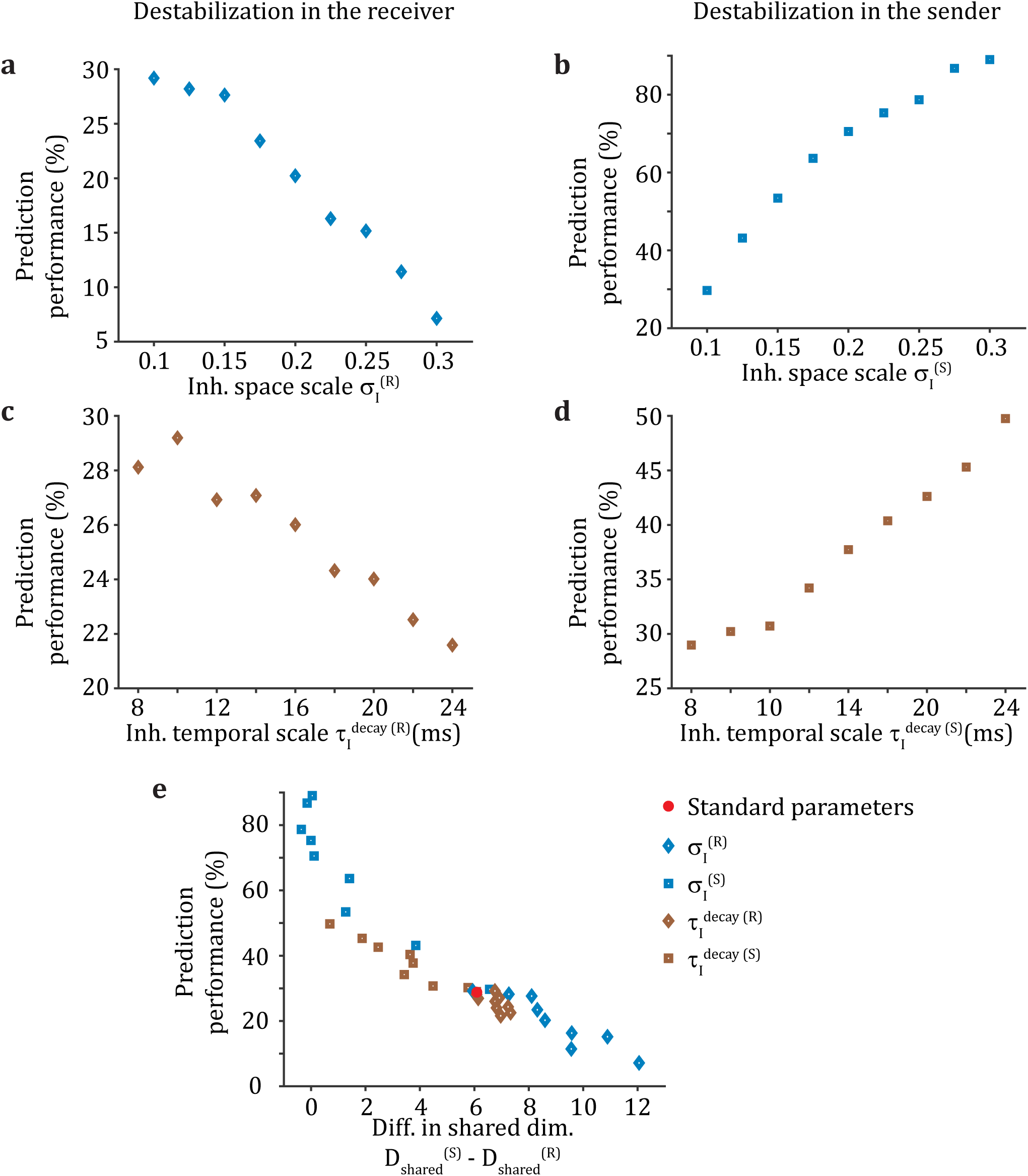
Inter brain-area communication strength depends on the origin of shared fluctuations. (**a**) When the E/I balance is spatially destabilized by broadening the inhibitory space scale *σ_I_*in the receiver, prediction performance of the communication subspace decreases. (**b**) When the E/I balance is spatially destabilized by increasing *σ_I_* in the sender, prediction performance of the communication subspace increases. (**c,d**) Similar to (a,b) when the E/I balance is temporally destabilized by increasing the inhibitory timescale *τ* ^decay^. (**e**) Prediction performance of the communication subspace is higher when there is a good match in shared dimensionality between sender and receiver (small difference in 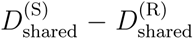) than when the sender is much higher dimensional than the receiver (large difference in 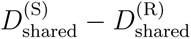). Neurons are sampled from a disc with radius 0.2; symbols are centered in the mean and include the SEM.

To provide a framework to organize these disparate results, we compute the difference in the shared dimensionality in the sender and receiver areas: 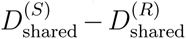. Strikingly, this difference determines the prediction performance across these varied datasets, irrespective of whether E/I balance is spatially or temporally destabilized in either the sender or the receiver (Fig. 4e). E/I destabilization in the sender through either broad or slow inhibition (Fig. 4e; blue and brown squares respectively) increases prediction performance compared to the standard network (Fig. 4e, red disc). By contrast, the same destabilization in the receiver decreases the performance compared to the standard network (Fig. 4e; blue and brown diamonds respectively). This organization of predictive performance by the comparison of sender versus receiver within-area dimensionality prompts a more complete analysis, which we present in the next section.

### A misalignment of shared variability is associated with poor communication between connected areas

Destabilization of E/I balance within a network often leads to low dimensional shared variability (Fig. 3). If the match in dimensionality of shared variability in the sender and receiver networks is all that is required for good communication (Fig. 4), then a combined E/I destabilization in the sender and receiver networks (so that both have low dimension) should yield a communication subspace with high prediction performance. If instead prediction performance is unexpectedly low, it would indicate that the matching in shared dimensionality is not the only requirement for faithful communication.

To test this hypothesis we compare the communication across two distinct sender – receiver networks. Both networks have the sender’s E/I balance destabilized through slow inhibition. In the first network the receiving population is kept in an intrinsically stable regime (Fig. 5a, top), while in the second network the receiver’s E/I balance is spatially destabilized (Fig. 5a, bottom). By design, both networks have a good match in dimensionality of the sender and receiver networks (Fig. S5). However, prediction performance of the communication subspace is substantially lower when different spatio-temporal patterns emerge in the sender and receiver networks individually, as compared to the case where the receiving area inherits its low dimensional nature from the sending area (Fig. 5b). Therefore, dimensionality matching is not a sufficient condition for faithful communication.

**Figure 5:**
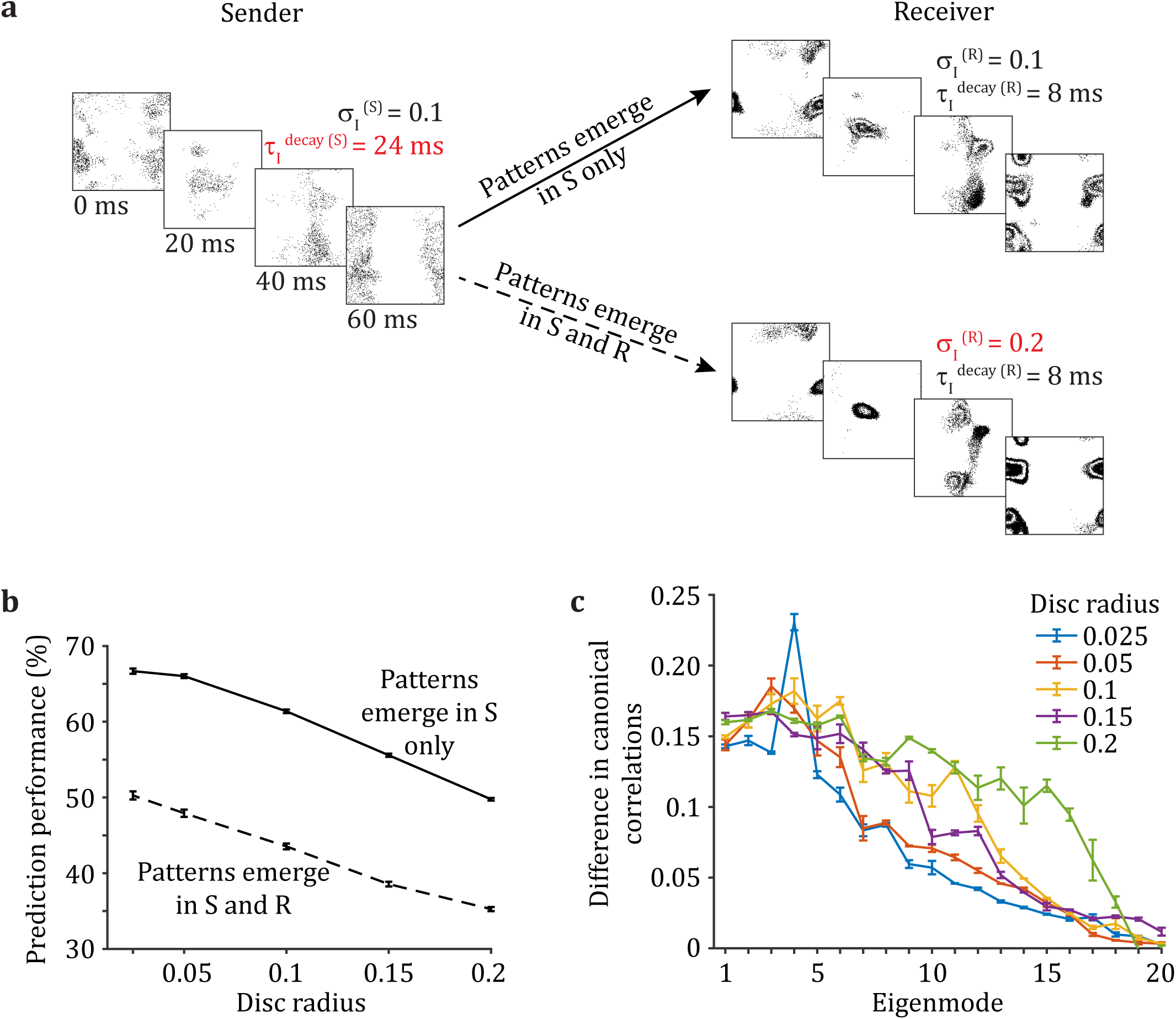
A misalignment of shared variability is associated with poor communication between connected areas. (**a**) The sender layer (S) is destabilized through a slower inhibitory timescale. The receiver layer (R) is either in the stable regime (top, patterns emerge in S only), or destabilized through broader recurrent inhibition (bottom, patterns emerge in S and R). (**b**) The prediction performance of the communication subspace between S and R is higher when only S is destabilized compared to when S and R are both destabilized. (**c**) Difference in canonical correlations between the case when only S is destabilized and the case when S and R are destabilized (see Fig. S5 for the canonical correlations in each case individually).

We hypothesize that a misalignment of the low dimensional manifolds of shared variability in the sender and receiver networks is the cause of disruption of their communication. We measure the alignment of shared variability in sender and receiver networks by computing their aligned canonical correlations. We conservatively select a dimensionality of the manifold of 20, which is higher than what was used in previous studies (Gallego et al., 2020). In our case, the value of the canonical correlation for each eigenmode indicates to what extent the given latent dimension can be well aligned between the sender and receiver networks through a linear transformation. We observe that the first eigenmodes can be well aligned and that the larger the spatial scale from which neurons are sampled, the better the alignment of their eigenmodes (Fig. S5). Importantly, we find that the canonical correlations between the sending and receiving populations are larger when only the sender network is destabilized, compared to when both sender and receiver networks are destabilized (Fig. 5c; this is true for all disc radii explored). Hence, a misalignment of the within brain-area shared fluctuations in sender and receiver populations can cause poor communication.

### Linear communication can be disrupted in two general ways

To summarize the results above, faithful communication as assessed by a linear measure is possible if shared dimensionality is similarly low in sender and receiver networks, and if the manifold of the shared variability of the receiver network can be well aligned to the one in the sending network. If the sender network displays high dimensional shared variability while the receiving population exhibits low dimensional shared variability, such as when spatio-temporal patterns of activity only emerge in the receiver network, communication is disrupted (Fig. 4a,c). Alternatively, if the shared variability in sending and receiving populations cannot be aligned through a linear transformation, for example when different spatio-temporal patterns emerge in both areas independently, the resulting communication is also poor (Fig. 5). We use these findings to investigate the underlying mechanisms by which linear communication between sender and receiver networks is disrupted.

If the within-area dimensionality of the sender and receiver networks is matched and their interaction is linear, then their communication is faithful (Fig. 6, top case). We outline two main hypothesized mechanisms by which linear communication can be disrupted. On one hand, because communication is assessed by a linear measure, if the mapping between sender and receiver activity is highly nonlinear, then communication will appear disrupted (Fig. 6, middle case). However, in this case the sender does effectively drive the receiver, yet this communication is blind to any linear metric. On the other hand, while the mapping of activity from sender to receiver could be linear, emergent dynamics in the receiver that is unrelated to the sender network activity may hinder communication (Fig. 6, bottom case). In this case, the emergent activity appears as ‘noise’ and corrupts the relation between the sender and receiver networks. In the remaining sections we show that the results presented above are classified into one of these two generic categories.

**Figure 6:**
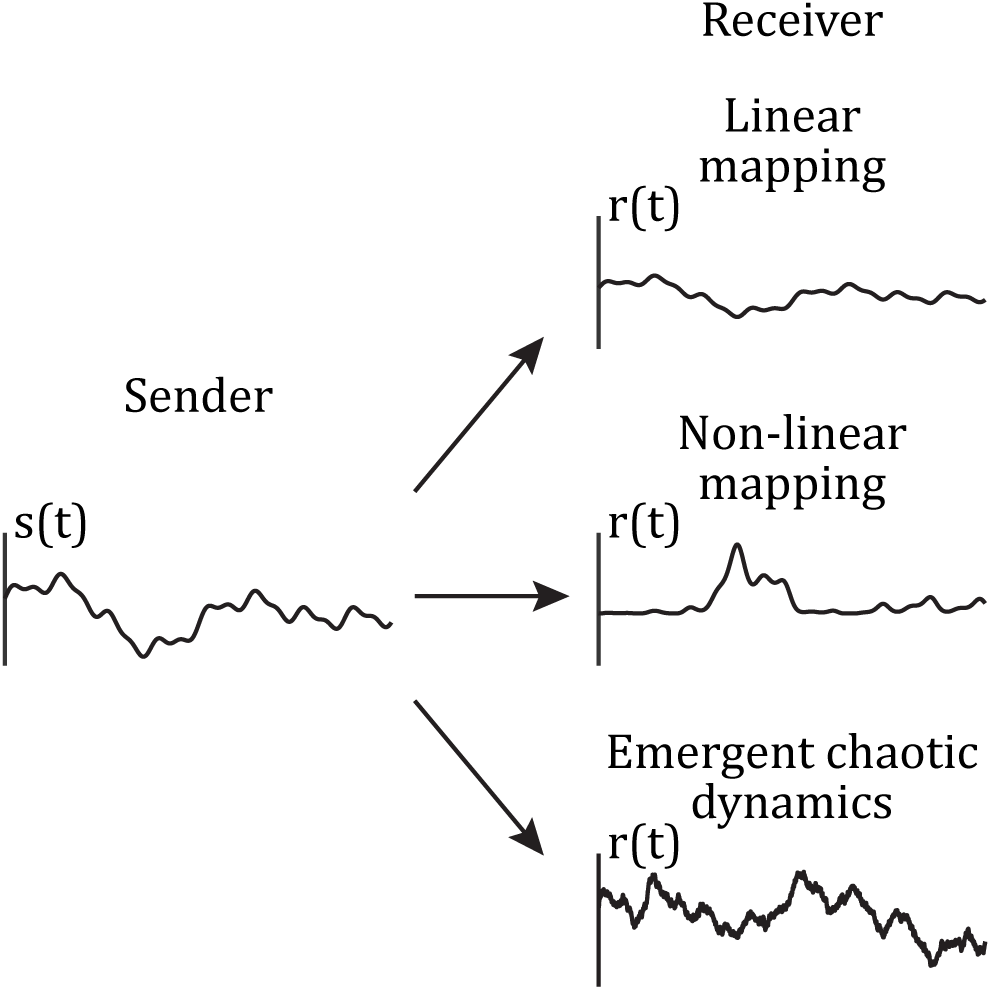
Linear communication can be disrupted in two general ways. For illustration purposes we represent the average activity in the sender and receiver as a scalar time series, *s*(*t*) and *r*(*t*), respectively. We set *s*(*t*) as a sum of six sine functions with amplitudes ranging over [0.05, 0.3], frequencies over [0.5, 10], and phases over [0, 2*π*]. If the mapping between sender and receiver is linear, such as the compression *r*(*t*) = 0.5*s*(*t*) (top receiver), then linear measures are suitable to uncover between-area communication. Rather, if the activity in the receiver is obtained through a nonlinear mapping of the activity in the sender, such as *r*(*t*) = −0.8*s*(*t*)(*s*(*t*) − 0.5)(*s*(*t*) − 1) (middle receiver), linear communication appears disrupted, even though the receiver is effectively driven by the sender. Finally, if dynamic fluctuations emerge in the receiver which are uncorrelated with the sender activity, then these fluctuations would act as ‘noise’ and degrade the linear communication between sender and receiver. In this illustration, we model these fluctuations as an additive Ornstein–Uhlenbeck process *n*(*t*) with *τ* = 10 s and *σ* = 0.05 and set *r*(*t*) = *s*(*t*) + *n*(*t*) (bottom receiver).

### Spatial but not temporal destabilization of the E/I balance yields a nonlinear mapping from sender to receiver

We determine whether the mapping of neuronal activity from sender to receiver networks remains linear despite destabilization of E/I balance in the receiver. We consider the activity of a destabilized output (receiver) network in response to an input (sender) population modeled by temporally homogeneous, independent Poisson spike trains. We compare the responses of the recurrently coupled output network constructed in two scenarios (Fig. 7).

**Figure 7:**
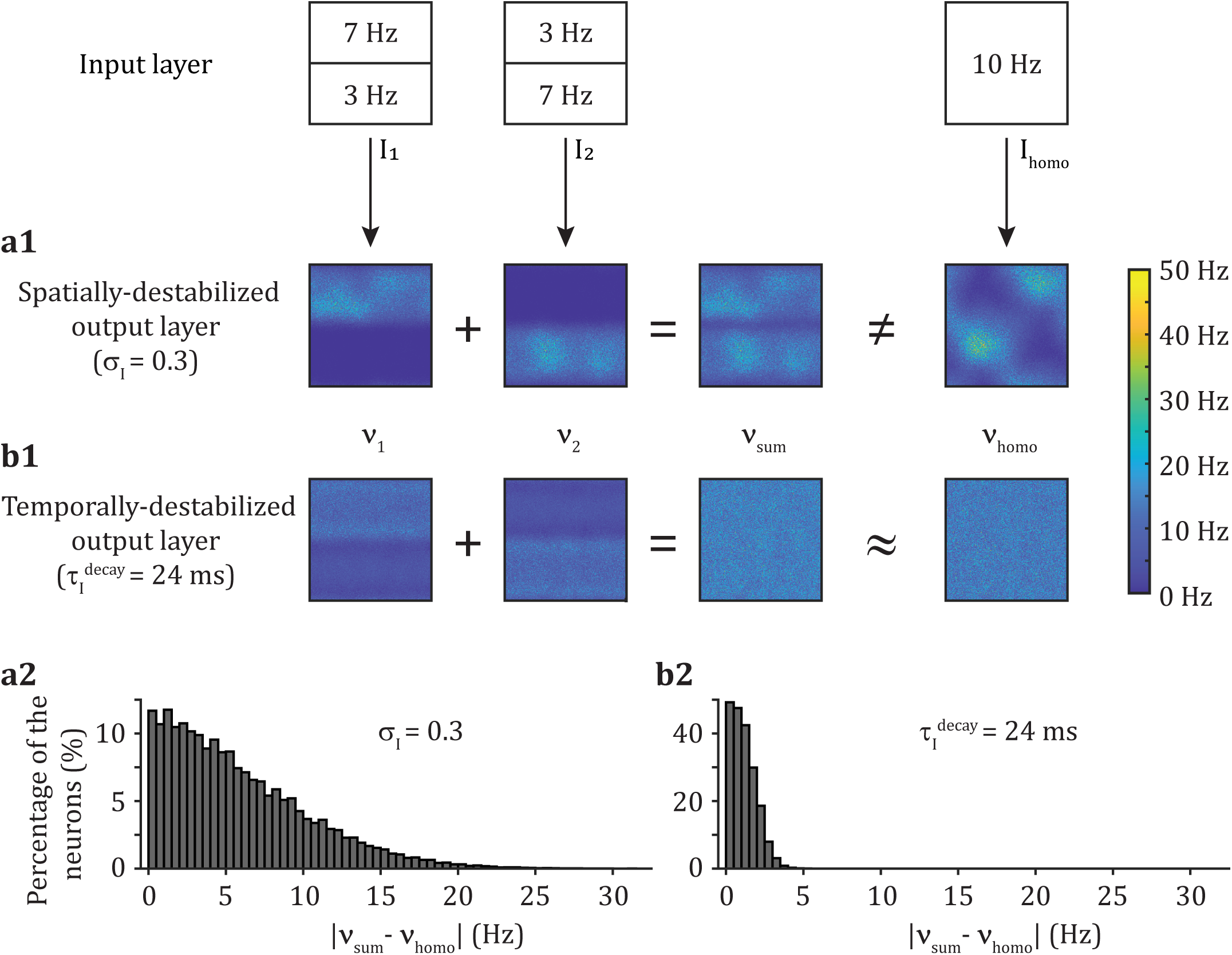
Spatial but not temporal destabilization of the E/I balance yields a nonlinear mapping from sender to receiver. The firing rate of the Poisson processes in the input layer is set to 7 Hz for the upper input neurons and 3 Hz for the lower neurons (*I*_1_), or the opposite (*I*_2_), or at 10 Hz for all input neurons (*I*_homo_). We compute the output layer response as ***ν***_1_, ***ν***_2_, and ***ν***_homo_ to these respective cases, and define ***ν***_sum_ = ***ν***_1_ + ***ν***_2_. (**a1**) When the output layer is spatially destabilized (*σ_I_* = 0.3), we have that ***ν***_sum_ */*= ***ν***_homo_. (**b1**) When the output layer is temporally destabilized (*τ_I_* ^decay^ = 24 ms), we have that ***ν***_sum_ *≈ **ν***_homo_. (**a2,b2**) Distribution of the absolute difference in firing rates between ***ν***_sum_ and ***ν***_homo_ for the case with a spatially-destabilized (a2) or a temporally-destabilized (b2) output layer. Each trial lasts 20s, from which we remove the first 1s, and estimate the firing rate of each neuron. The panels show the results for one trial. Over 30 trials, the mean absolute difference in firing rate is (*µ* std): 5.3 4.2 Hz for the spatially-destabilized network, and 1.1 0.8 Hz for the temporally-destabilized network.

The first scenario provides spatially heterogeneous input *I*_1_ where the neurons located in the upper and lower halves of the spatial grid receive inputs with firing rates of 7 Hz or 3 Hz, respectively. We record the (vectored) output neuron firing rates ***ν***_1_ in response to this input. We repeat this experiment but with the input *I*_2_ where the upper and lower half input rates are switched, and we record response rates as ***ν***_2_. We construct the response to the summed input *I*_1_ + *I*_2_ to be ***ν***_sum_ = ***ν***_1_ + ***ν***_2_. The second scenario provides spatially homogeneous inputs *I*_homo_ = *I*_1_ + *I*_2_ so that all neurons receive inputs with a firing rate of 10 Hz, and we record the response ***ν***_homo_. A linear mapping between input and output would predict that ***ν***_sum_ = ***ν***_homo_.

A spatial destabilization of the E/I balance in the output layer shows a summed response activity that is different from the activity in response to the homogeneous input (Fig. 7a1). Indeed, the lateral inhibition in the output layer patterns the response to the homogeneous input which disagrees with a simple linear mixing of the responses. Beyond a simple visual inspection of the spatial distribution of firing rates, we compute for every neuron *j* in the output network the comparison |*ν*_sum*,j*_ − *ν*_homo*,j*_| (Fig. 7a2). The absolute difference is quite large, indicating a complete reshuffling of individual neuron firing rates between scenarios, implying that across the output network we have that ***ν***_sum_ ≠ ***ν***_homo_. Thus, a spatially destabilized E/I balance imparts a clear nonlinear mapping between input and output spiking activity.

By contrast, a temporally destabilized E/I balance in the output network produces no obvious change in spatial distribution of firing rates (Fig. 7b1). When individual neurons are compared across scenarios, we observe only a slight change between *ν*_sum*,j*_ and *ν*_homo*,j*_ (Fig. 7b2), so that we have across the output network ***ν***_sum_ ≈ ***ν***_homo_. Hence, a linear mapping of activity is qualitatively maintained despite temporal destabilization of the output layer.

These results suggest that when the E/I balance in the receiver is spatially destabilized a nonlinear mapping between sender and receiver activity (Fig. 7a) contributes to the disruption of linearly measured communication (Fig. 4a). However, while temporal destabilization in the receiver disrupts linear communication (Fig. 4c) it does maintain a sender to receiver linear mapping (Fig. 7b). Therefore, in the next section we explore the hypothesis that temporal destabilization induces emergent dynamics in the receiver network which corrupt the communication between the sender and receiver.

### Temporal but not spatial destabilization of the E/I balance yields macroscopic emergent dynamics in the receiver

Destabilization of E/I balance in the receiver network, through either broad or slow inhibition, produces rich, spatio-temporal patterns of spiking activity (Fig. 2). However, it is unclear whether this activity corrupts communication between the sender and receiver networks. To determine this, we generate a single (frozen) realization of input spike trains from the input layer and record the resulting spike train activity from the sender layer. To this frozen sender input we compare two response trials in the receiving layer, where the only difference between them is the initial membrane voltage of the excitatory neurons in the receiving network (Fig. 8a,b). If the patterns of spiking activity in the receiver network differ significantly between trials despite the sender activity being frozen, then this would indicate an emergent, chaotic dynamic owing to complex recurrent interactions (Van Vreeswijk and Sompolinsky, 1996; London et al., 2010; Monteforte and Wolf, 2012). In this case the differing receiver network responses across trials would act as ‘noise’ that corrupts the communication between sender and receiver.

**Figure 8:**
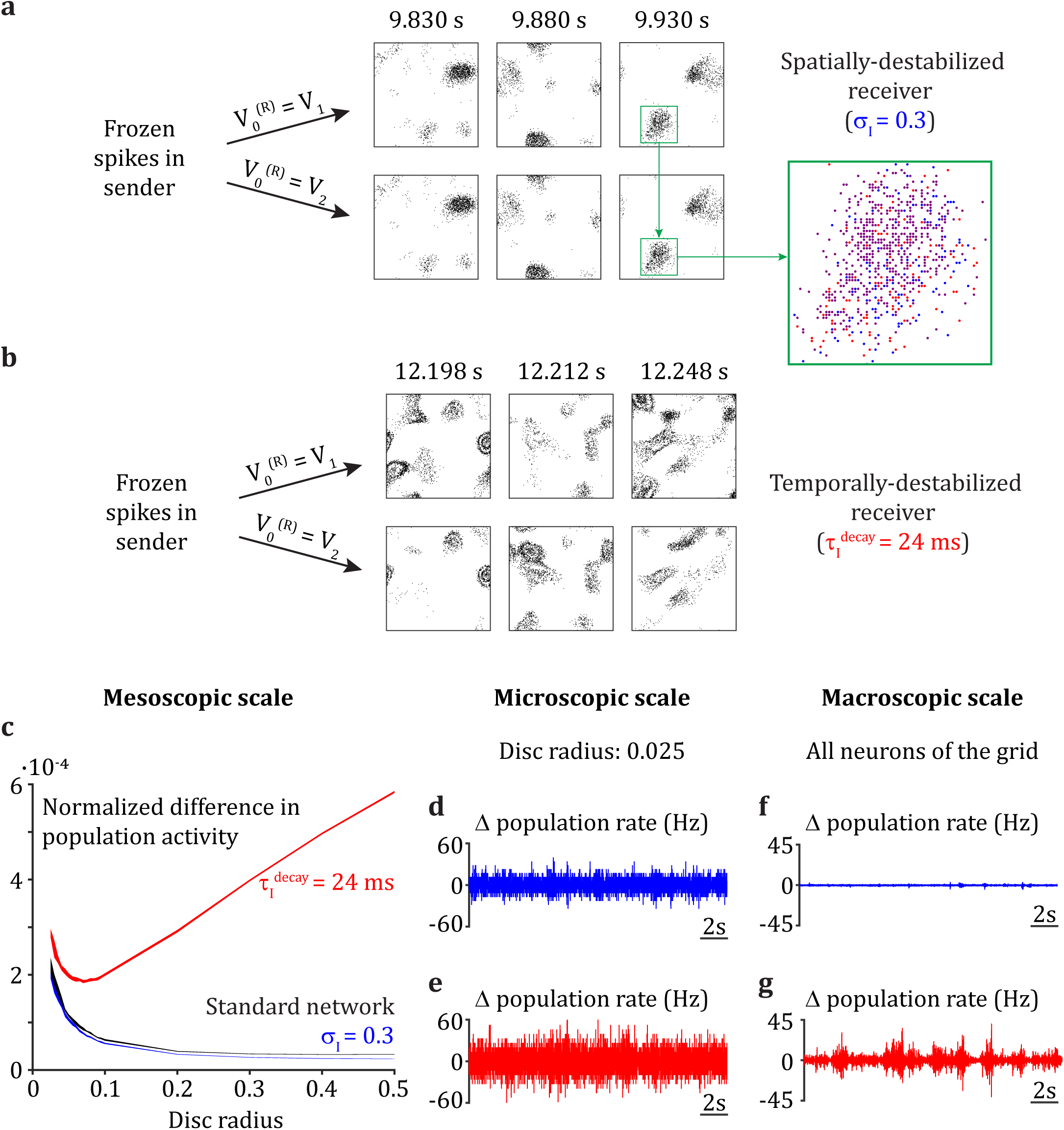
Temporal but not spatial destabilization of the E/I balance yields macroscopic emergent dynamics in the receiver. (**a,b**) A single realization of spiking activity is generated in the input layer and propagated to the sender (S). The membrane potential of neuron *j* in the receiver (R) is set to one of two different initial conditions: *V_j_* ^1^ or *V _j_*^2^ (randomly generated for each neuron). Spike time raster plots (Δ*t* = 2 ms) at three time points are presented for the network with *σ_I_*^(*R*)^ = 0.3 (a), and for the network with *τ_I_* ^decay(*R*)^ = 24 ms (b) (*σ_I_*^(*S*)^ = 0.1 and *τ _I_*^decay(*S*)^ = 8 ms in both cases). Raster plot zoom in: blue dots are spike times for the first trial, red dots are spike times for the second trial and purple dots are overlapping spike times of the first and second trials. (**c**) Normalized difference in population activity for neurons sampled from a disc of radius 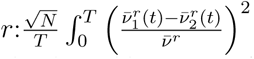, with *N* the number of neurons within the disc, *T* the length of the trials (20s), 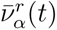 for *α* ∈ 1, 2 being the average firing rate over all neurons of the disc for realization 1 and 2, and *ν̄^r^* the average firing rate over all neurons within the disc from both trials and all time points. (**d,e**) Difference in the average firing rate over all neurons sampled from a random disc with radius 0.025 (96 neurons) between the two trials with different initial conditions as a function of time; for the spatially-destabilized case (d), and the temporally-destabilized case (e). (**f,g**) Same as (d,e), but for the average firing rate over all neurons of the grid (40,000).

When E/I balance in the receiver network is spatially destabilized, the difference in initial membrane potential conditions does not affect the overall macroscopic (aggregated) patterns of spiking activity across the two trials (Fig. 8a). By contrast, when the receiver is temporally destabilized, we observe significant discrepancy in the macroscopic spatio-temporal patterns between the two trials (Fig. 8b). Therefore, these initial results support our hypothesis that temporal E/I destabilization induces emergence of chaotic dynamics at the population-wide scale which is the source of linear communication disruption. However, even during spatial E/I destabilization, while the macroscopic (population-wide) network activity is reliable across trials, the microscopic activity is nevertheless unreliable. The exact spike train sequences across the network differ significantly between the two trials (Fig. 8a, zoom in), so that any observer of a single neuron would easily distinguish the trials. These results agree with previous studies where a weak perturbation to network activity yields only a transient change in firing rate but a long-lasting trial-to-trial decorrelation of spike sequences in spatially disordered balanced networks (London et al., 2010; Monteforte and Wolf, 2012). The differential spatial scale (micro versus macro) of unreliability in spatially and temporally E/I destabilized networks prompts a more detailed analysis.

We consider the trial-to-trial reliability of the spiking activity of neurons aggregated over a spatial disc of radius *r*. Across a trial we sum the instantaneous spiking activity of the neurons within the disc, *ν^r^*(*t*). We integrate the normalized difference of *ν^r^*(*t*) between trials as a measure of unreliability (Fig. 8c). Spatial destabilization only yields microscopic unreliability, as observed for discs of small radii (Fig. 8d,f), similar to the standard network in the stable regime (Fig. 8c, compare blue and black curves). In contrast, a temporal destabilization induces macroscopic unreliability in addition to microscopic unreliability (Fig. 8c, red curve; e,g). These results extend our understanding of rich spiking neuronal dynamics in networks with biologically-constrained architecture and are consistent with the observation of macroscopic chaotic dynamics upon temporal – but not spatial – destabilization of the E/I balance in rate networks (Mosheiff et al., 2023).

## Discussion

The mechanisms that produce low dimensional shared variability across a neuronal population can be organized into two broad categories. First, shared variability of the neurons in a brain region may be inherited (in part) from connecting brain areas (Wimmer et al., 2015; Gómez-Laberge et al., 2016; Semedo et al., 2019). Second, shared variability may be an emergent property of a brain area, owing to local, complex recurrent interactions between neurons (Darshan et al., 2018; Landau and Sompolinsky, 2018; Mastrogiuseppe and Ostojic, 2018; Huang et al., 2019). Population recordings restricted to a single brain area cannot easily disentangle the contributions of each mechanic to the total shared variability. Rather, multi-area brain recordings will be needed to expose these separate mechanisms (Urai et al., 2022). In our study we explored signatures of inherited or emergent shared variability within a brain area by measuring the (linear) communication between distinct, but connected, brain areas.

A suitable modeling framework to investigate the interplay between inheritance and emergence of neuronal variability in brain circuits has only recently become available. At one extreme the inheritance of neuronal fluctuations has been extensively studied through the analysis of activity propagation in feedforward networks (Abeles, 1991; Diesmann et al., 1999; Reyes, 2003; Rosenbaum et al., 2011). However, since those circuits explicitly lacked within layer recurrent connections, they could not model the emergence of complex population dynamics. At the other extreme, within-population recurrent, yet unstructured, excitatory and inhibitory connections were introduced to model asynchronous activity reminiscent of the baseline state of cortical variability (Van Vreeswijk and Sompolinsky, 1998; Amit and Brunel, 1997; Renart et al., 2010). However, these networks did not produce rich, low dimensional fluctuations shared across the population. Over the past few years novel modeling frameworks have included structure to the within-population recurrent wiring that permits low dimensional fluctuations to intrinsically emerge within the network. Different approaches have been taken. On one hand, forcing a low rank structure of the recurrent connectivity matrix yields low dimensional activity, revealing a strong relationship between structure and dynamics (Mastrogiuseppe and Ostojic, 2018; Landau and Sompolinsky, 2018). On the other hand, low dimensional shared fluctuations can emerge due to a destabilization of E/I balance despite a connectivity matrix with high rank (Rosenbaum et al., 2017; Darshan et al., 2018; Huang et al., 2019). In our work, we leverage those last modeling frameworks to investigate how the emergence and inheritance of low dimensional neuronal fluctuations affect between-area interactions in well controlled settings.

We assess the interaction between a sender network and a receiver network through a linear communication measure. Brain recordings are believed to frequently operate in a linear regime, in particular in early sensory areas (Stringer et al., 2021). Besides, linear methods are routinely used for the analysis of neuronal dynamics and in many cases have proven successful. At the single population level, they have unraveled stable low-dimensional manifolds in working memory networks (Murray et al., 2017) as well as in motor areas (Jiang et al., 2020). In addition, the suppression of rich spatio-temporal dynamics has been shown to increase the amount of linear Fisher information that is propagated down successive layers (Huang et al., 2022). However, cognition arises through the interaction of distributed brain regions. Recent technological advances have allowed a glimpse into distributed processing through simultaneous recordings from distinct neuronal populations, allowing us to answer the question of how much of the dynamics in a population can be explained by the activity of another recorded brain area (Yu et al., 2019; Musall et al., 2019). At this multiple populations level, linear methods have also been critical. They have exhibited selectivity (Kaufman et al., 2014) and a low dimensional subspace (Semedo et al., 2019; Srinath et al., 2021) in the communication between brain areas. Our work shows that those methods provide a useful lens to investigate the inheritance of neuronal fluctuations by a receiver displaying linear mapping, in particular in low dimensional inheritance settings.

However, when the receiving area is in a nonlinear regime where complex spatio-temporal dynamics intrinsically emerge, we show that between-area communication cannot be properly assessed by linear measures. Our results support a recent study revealing that when a network exhibits strong pairwise correlations, reminiscent of low-dimensional pattern formation, connectivity inference is biased towards an excess of connectivity between highly correlated neurons (Das and Fiete, 2020). Indeed, linear inference methods are only appropriate when neuronal dynamics operate in a linear regime, where activity is high-dimensional and unstructured (Das and Fiete, 2020). Even though early sensory areas are believed to mostly operate in a linear regime to faithfully encode sensory inputs, brain areas involved in higher-level cognition exhibit low dimensional spatio-temporal dynamics (Wang, 2002; Mante et al., 2013; Lara et al., 2018; Chen et al., 2021). Besides, it has recently been shown that microscopic irregularity can subsist even in the presence of macroscopic fluctuations (Pyle and Rosenbaum, 2017; Darshan et al., 2018). Our results indicate that even when the receiver network is in the pattern-forming regime due to a spatial destabilization, it is nevertheless reliably driven by the sender network, as reflected by a lack of macroscopic chaos. However, a nonlinearity in the sender – receiver mapping makes linear methods of communication blind to this interaction. Alternatively, emergence of unreliable macroscopic spiking dynamics due to temporal destabilization also impair linear measures of communication even though the interaction is effectively linear. Therefore, novel nonlinear methods, as well as ways to characterize emergent fluctuations, will have to be developed to accurately assess communication involving brain areas with more complex dynamics, such as those involved in higher-level brain regions associated with cognition.

The simplicity of our modeling framework is critical to thoroughly study the interplay between inheritance and emergence of neuronal fluctuations in tightly controlled settings. The use of feedforwardly connected distinct sender and receiver networks, each only involving local recurrent connections, allows us to observe the differential effects of low dimensional shared fluctuations on between-area communication depending on their well defined origin. However, brain circuits show recurrent architecture spanning a wide range of spatial scales. Furthermore, cognition is believed to arise from distributed computational processes. Even in early sensory areas historically thought of as mostly feedforward, the importance of feedback interactions in sensory processing has started to be exposed (Semedo et al., 2022). Therefore, a natural extension of our work will be the implementation of more complex circuit architectures, starting by the introduction of feedback connections, to provide novel insights into the mechanistic interplay of inheritance and emergence of shared fluctuation across spatial scales.

## Methods

### Network structure

A three-layer spiking network model is implemented similar to previous work (Huang et al., 2019). The input layer consists of 2,500 excitatory neurons whose spikes are taken from independent homogeneous (space and time) Poisson processes with a uniform rate of 10 Hz. The sender and receiver layers each consist of 40,000 excitatory (E) and 10,000 inhibitory (I) neurons which are arranged on a unit square domain Γ = [0, 1] *×* [0, 1] with periodic boundary conditions. The probability of connection between a presynaptic neuron belonging to class *β ∈ {*E,I*}* located at position *y⃗* = (*y*_1_*, y*_2_) and a postsynaptic neuron belonging to class *α ∈ {*E,I*}* located at position *x⃗* = (*x*_1_*, x*_2_) depends on their pairwise distance measured periodically on Γ:

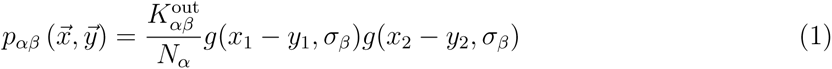

where *K_αβ_*^out^ is the out-degree, so 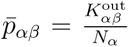 is the mean connection probability, and *g*(*u, σ*) is a wrapped Gaussian distribution:

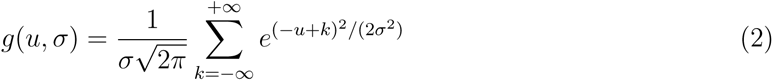

Excitatory feedforward connections between layers and recurrent excitatory and inhibitory connections within layers are spatially distributed according to a Gaussian with width *σ*_ffwd_, *σ_E_* and *σ_I_*respectively.

### Neuronal dynamics

Excitatory and inhibitory neurons in sender and receiver networks are modeled as conductance-based exponential integrate-and-fire neurons:

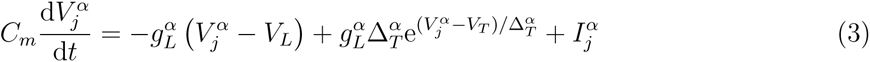

with *α* E, I. A neuron spikes when its membrane voltage *V_j_^α^* reaches the spiking threshold *V*_th_. Then its membrane voltage is reset at *V*_reset_ for a refractory period *t*_ref_*^α^*. The total current received by neuron *j* belonging to class *α*, *I_j_^α^*(*t*), is given by the summation of feedforward and recurrent input:

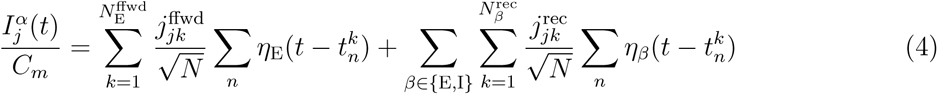

with *N* = *N*_E_ + *N*_I_ the total number of neurons within the layer of interest. The postsynaptic current, *η_β_*(*t*), is induced by presynaptic spiking and depends on the class (*β* E, I) of the presynaptic neuron. Assuming a single presynaptic spike at time *t* = 0, it is given by the difference of two exponentials with rise timescale *τ_β_*^rise^ and decay timescale *τ_β_*^decay^:

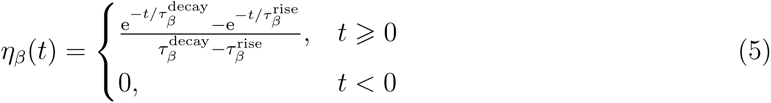

Equations are numerically integrated using the forward Euler method with a timestep of 0.05 ms. Instantaneous firing rates were computed from the spike counts in non-overlapping 50 ms bins. Unless specified otherwise, all neuronal and connectivity parameters are identical in the sender and the receiver layers (Table 1).

### Shared covariance matrix and within-area shared dimensionality

Shared covariance of within-area neuronal activity *x⃗* is assessed through factor analysis (FA):

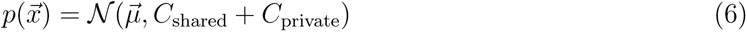

where *C*_private_ is a diagonal matrix whose elements represent the individual neuronal variances and *C*_shared_ represents the shared component of the full covariance matrix (Everitt, 1984; Yu et al., 2009). Singular value decomposition is applied to *C*_shared_:

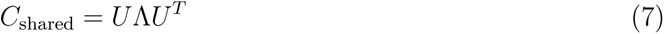

where the columns of *U* are the eigenvectors and the elements of the diagonal matrix Λ, *λ_i_*, are the associated eigenvalues ordered from larger to smaller.

The dimensionality of the shared covariance matrix is estimated using the effective dimension (sometimes referred to as the participation ratio) (Mazzucato et al., 2016; Litwin-Kumar et al., 2017):

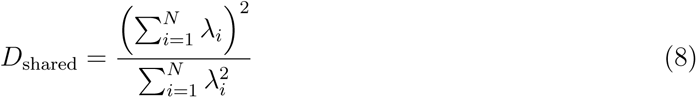

### Between brain-area communication

To assess communication between sender and receiver networks, we use a recently developed communication subspace measure based on reduced-rank regression (Semedo et al., 2019):

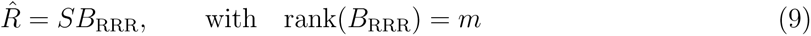

with *S* the activity in the sender network and *R̂* the estimated activity in the receiver network (both of size [*T × K*]). *T* is the total number of timepoints and *K* the number of sampled neurons (we set *K* = 50 in both the sender and the receiver). The reduced-rank regression matrix, *B*_RRR_, is a low-rank approximation of the ordinary least-squares solution (Semedo et al., 2019):

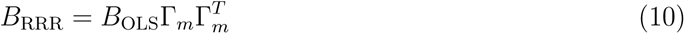

with *B*_OLS_= *S^T^ S ^−^*^1^ *S^T^ R* a full rank regression matrix. Hence, Γ_m_ is a matrix whose columns are the *m* first eigenvectors of the covariance matrix of the receiver activity estimated using the ordinary least-squares solution: (*SB*_OLS_)*^T^* (*SB*_OLS_) = ΓΛΓ*^T^*. The dimensions in activity *S* that are most predictive of activity *R* according to the communication subspace measure are called “predictive dimensions”. They are the *m* columns of *B*_OLS_Γ*_m_*.

We define the prediction performance as the R-Square value (RSS: residual sum of squares, TSS: total sum of demeaned squares):

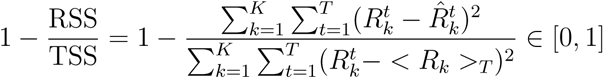

where 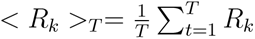. In the results, we multiply the prediction performance by 100 to get percentages.

For each network parameter set, we compute prediction performance of a communication sub- space with rank *m*, *m* 0, 1*, …, K*. We define the optimal rank *m^∗^* as the smallest number of dimensions for which the prediction performance is within one SEM of the peak performance over 20 cross-validation folds. In the results, we report the average prediction performance using *B*_RRR_ with optimal rank *m^∗^* over 20 cross-validation folds.

### Alignment of shared variability

To align the simultaneously recorded latent dynamics in the sending and receiving populations, we use a method based on canonical correlation analysis which has recently been used to align within-area latent dynamics over several days where a different number of neurons with different identities were recorded (Gallego et al., 2020). We know from equation (7) that the singular value decomposition of the shared covariance matrix is given by: *C*_shared_ = *U* Λ*U^T^*. Hence the columns of *U* are the eigenvectors of *C*_shared_, ordered from the one which explains the most shared variance to the one that explains the least. We keep *m* = 20 shared latent variables, as they are sufficient to explain most of the shared variance. Therefore, we only keep the first *m* columns of *U* : *U_m_* of size [*K × m*] with *K* = 50 the number of sampled neurons. For the sender and receiver networks separately, we project the activity of the *K* sampled neurons onto the *m* shared latent variables: *L_k_* = *XU_m_*. We thus obtain two [*T × m*] matrices of latent dynamics *L_k_*, where *T* is the number of timepoints, *k ∈ {S, R}* and *X* is the corresponding activity in the sender (S), or in the receiver (R), of size [*T × K*]. Then we compute the QR decomposition of the shared latent dynamics: *L_k_* = *Q_k_R_k_*. The singular value decomposition of the covariance of *Q_S_* with *Q_R_* is given by: *Q_S_^T^ Q_R_* = *ŨS̃Ṽ^T^*. Canonical correlation analysis finds new latent directions to maximize the pairwise correlations between the sending and receiving populations. The projection of the shared latent dynamics onto these new latent directions is given by: *L̃_k_* = *L_k_M_k_*, where *M_S_* = *R_S_^−^*^1^*Ũ* and *M_R_* = *R_R_^−^*^1^*Ṽ*. Finally, the aligned canonical correlations are given by the pairwise correlations between the aligned shared latent dynamics: *L̃*_S_*^T^ L̃_R_* = *Ũ^T^ Q_S_^T^ Q_R_Ṽ* = *S̃*. They are the elements of the diagonal matrix *S̃*, which are ordered from larger to lower value.

### Code sharing

All associated simulation and analysis code is freely available at https://github.com/ogozel/neuronal_dynamics

## Acknowledgements

This work was supported by grants from the Simons Foundation Collaboration on the Global Brain, the National Institutes of Health Grants (1U19NS107613-01 and R01 EB026953), and the Vannevar Bush Faculty Fellowship (N00014-18-1-2002).

---

**Figure S1:**
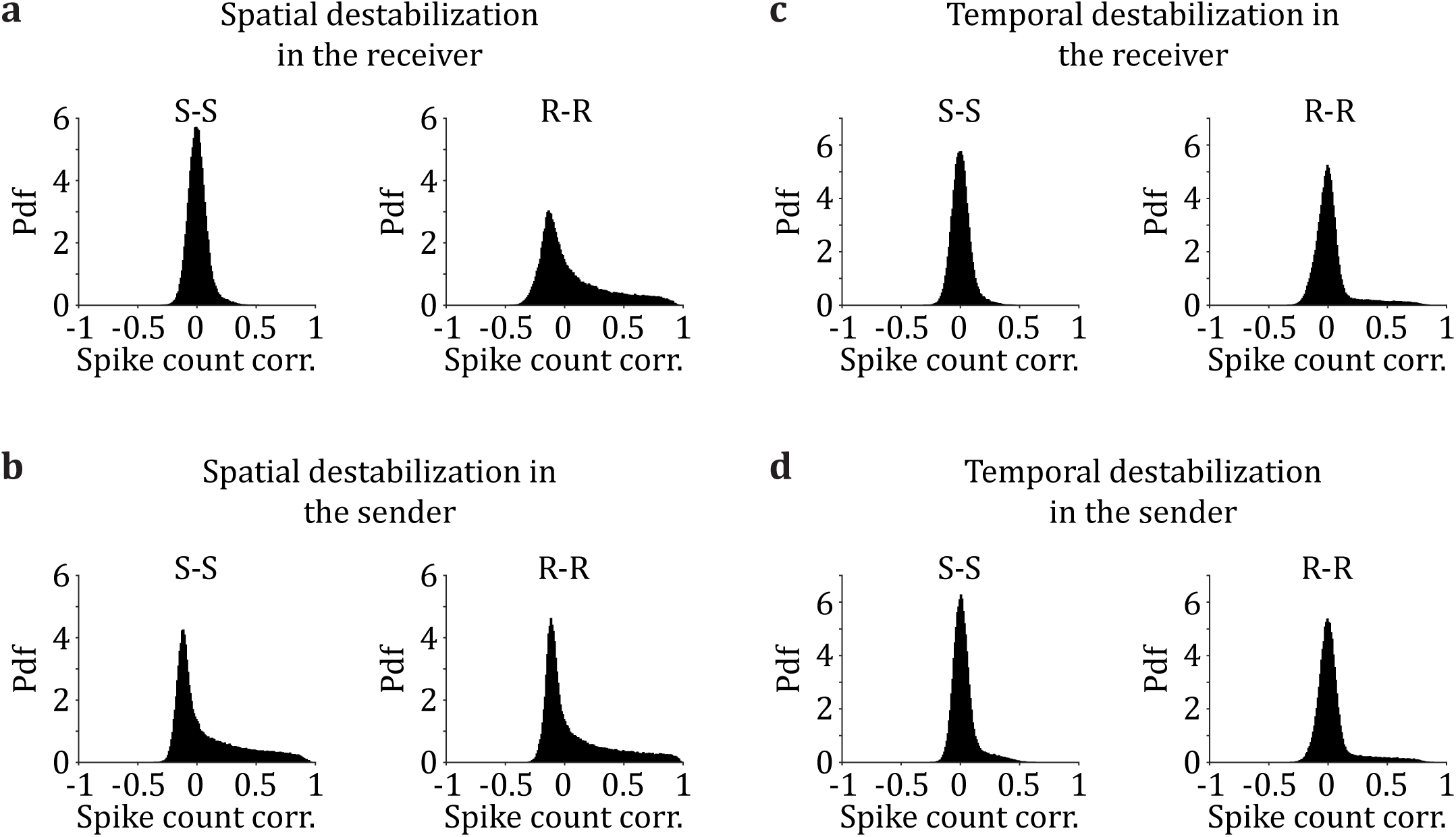
Within-area spike count correlations are differentially affected by the type of E/I balance destabilization. (a) Probability density functions (pdfs) of the pairwise spike count correlations for neuron pairs within the sender area, S (left, mean of the distribution: *µ* = 0.0045), and within the receiver area, R (right, *µ* = 0.0658) when the E/I balance is destabilized spatially in the receiver (*σ*_I_^(R)^ = 0.3). (b) Pdfs of the pairwise spike count correlations for neuron pairs within-area S (*µ* = 0.0839), and within-area R (*µ* = 0.0866) when the E/I balance is destabilized spatially in the sender (*σ*^(S)^ = 0.3). (c) Pdfs of the pairwise spike count correlations for neuron pairs within S (*µ* = 0.0043), and within R (right, *µ* = 0.0237) when the E/I balance is destabilized temporally in the receiver (*τ_I_*^decay (R)^ = 24 ms). (**d**) Pdfs of the pairwise spike count correlations for neuron pairs within-area S (*µ* = 0.0269), and within-area R (*µ* = 0.0394) when the E/I balance is destabilized temporally in the sender (*τ* ^decay^ ^(S)^ = 24 ms).

**Figure S2:**
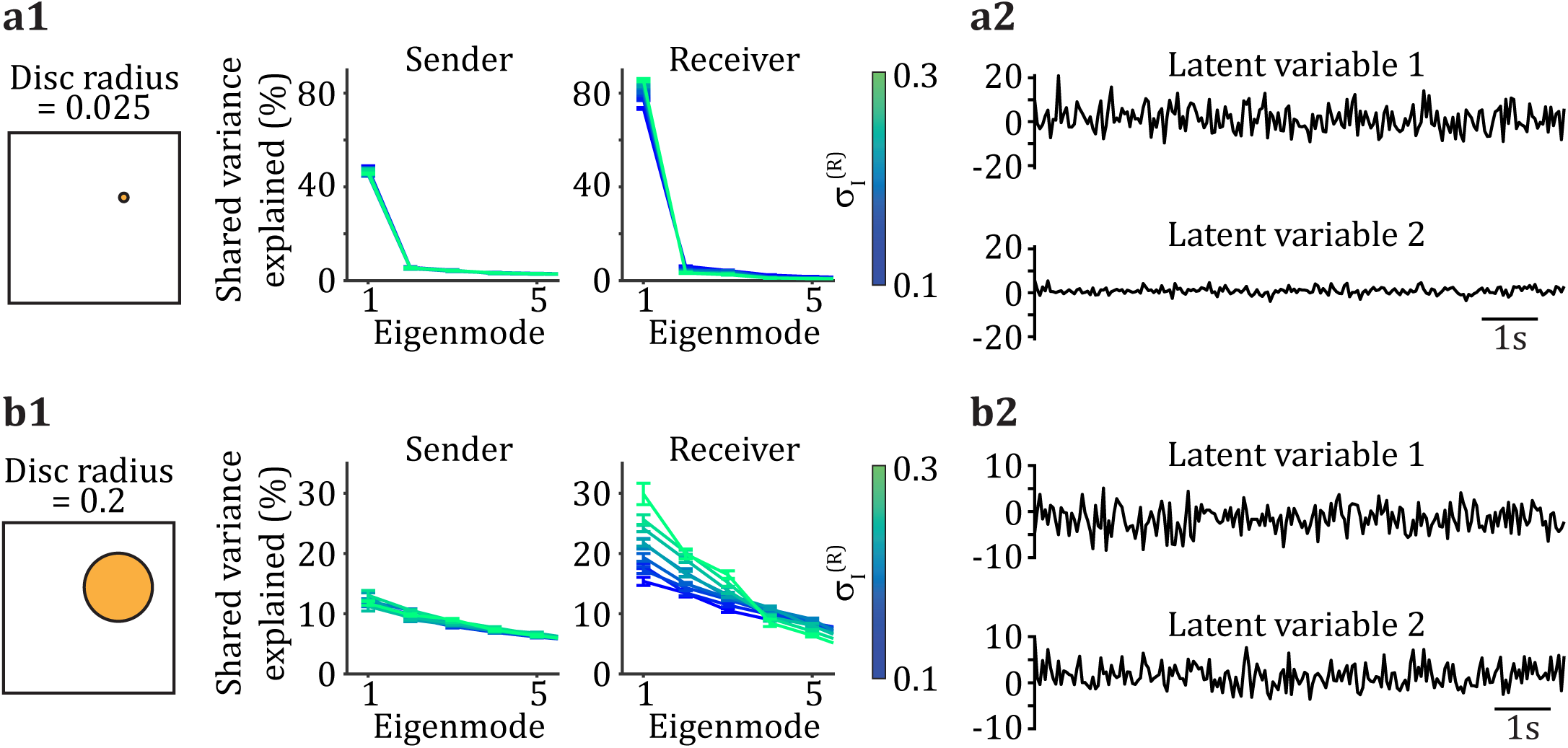
The structure of within-area shared fluctuations is spatial-scale dependent. (**a1**) Shared variance explained by the first five eigenmodes within the sender network (S) and the receiver network (R) when modifying *σ_I_*in R. Neurons are randomly sampled from a small disc (disc radius = 0.025). (**a2**) Projection of 10 s of the receiving population activity on the first two shared eigenvectors to obtain the first two shared latent variables. Neurons are sampled from a small disc in the standard network (*σ_I_*^(*S*)^ = σ_I_^(*R*)^ = 0.1). (**b1,b2**) Same as (a1,a2), except that neurons are sampled from a large disc (disc radius = 0.2).

**Figure S3:**
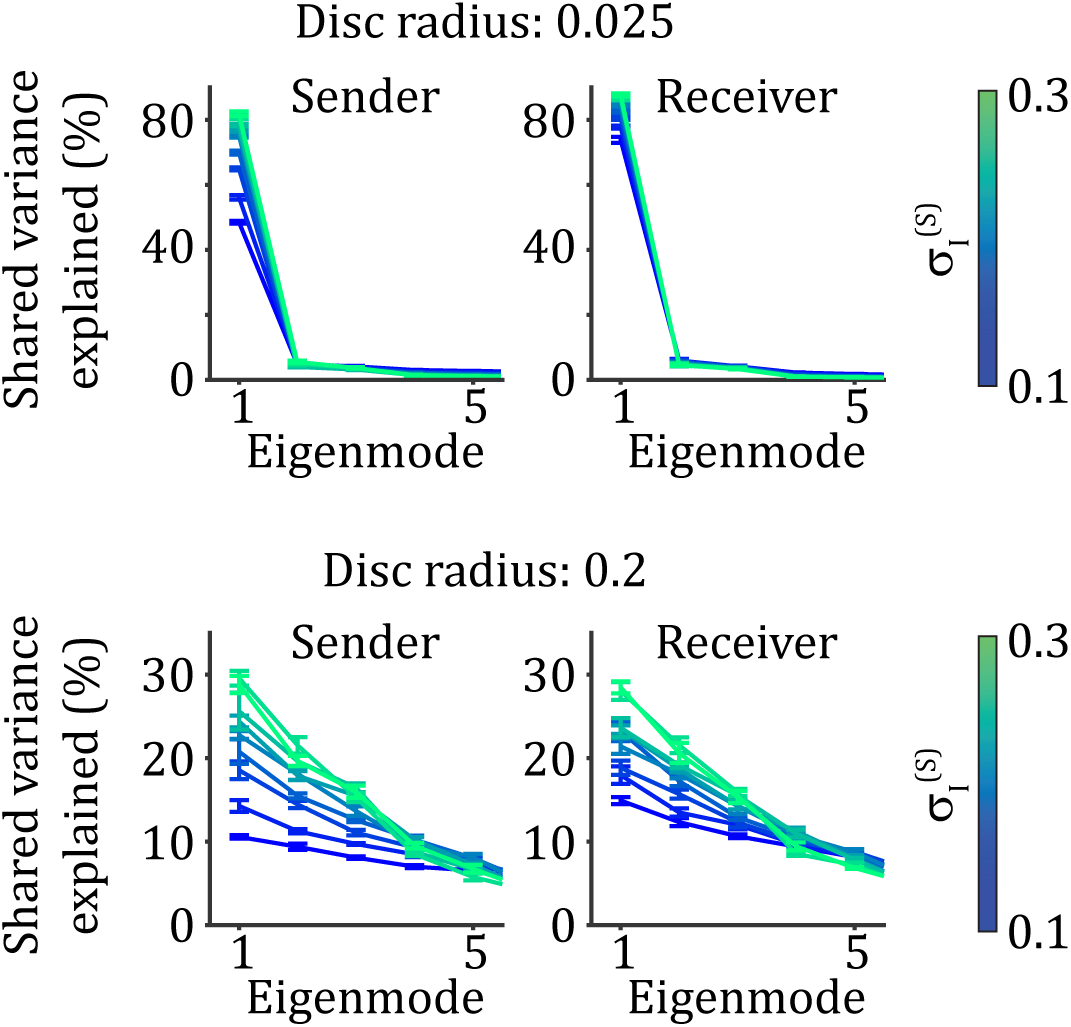
The structure of within-area shared fluctuations is spatial-scale dependent. Shared variance explained by the first five eigenmodes within sender network (S, left) and within receiver network (R, right) when modifying *σ_I_* in S. Neurons are randomly sampled from a small disc (top) or from a large disc (bottom).

**Figure S4:**
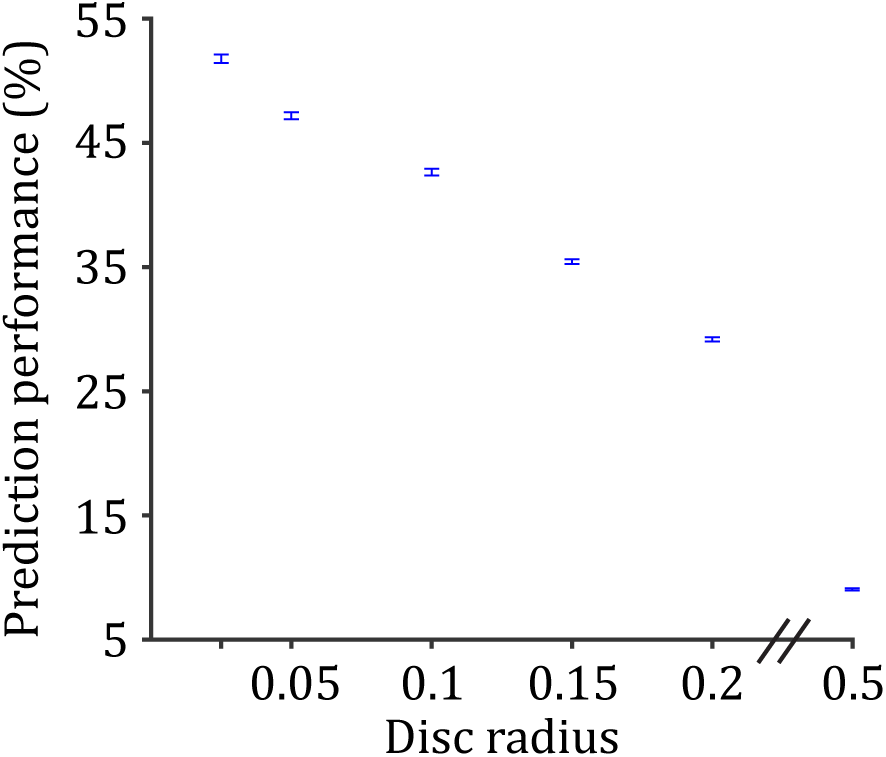
Dependence on the size of the spatial domain from which neurons are sampled on prediction performance of the communication subspace. Scenario with the standard network parameters (*σ_I_*^(*S*)^ = *σ_I_*^(*R*)^ = 0.1).

**Figure S5:**
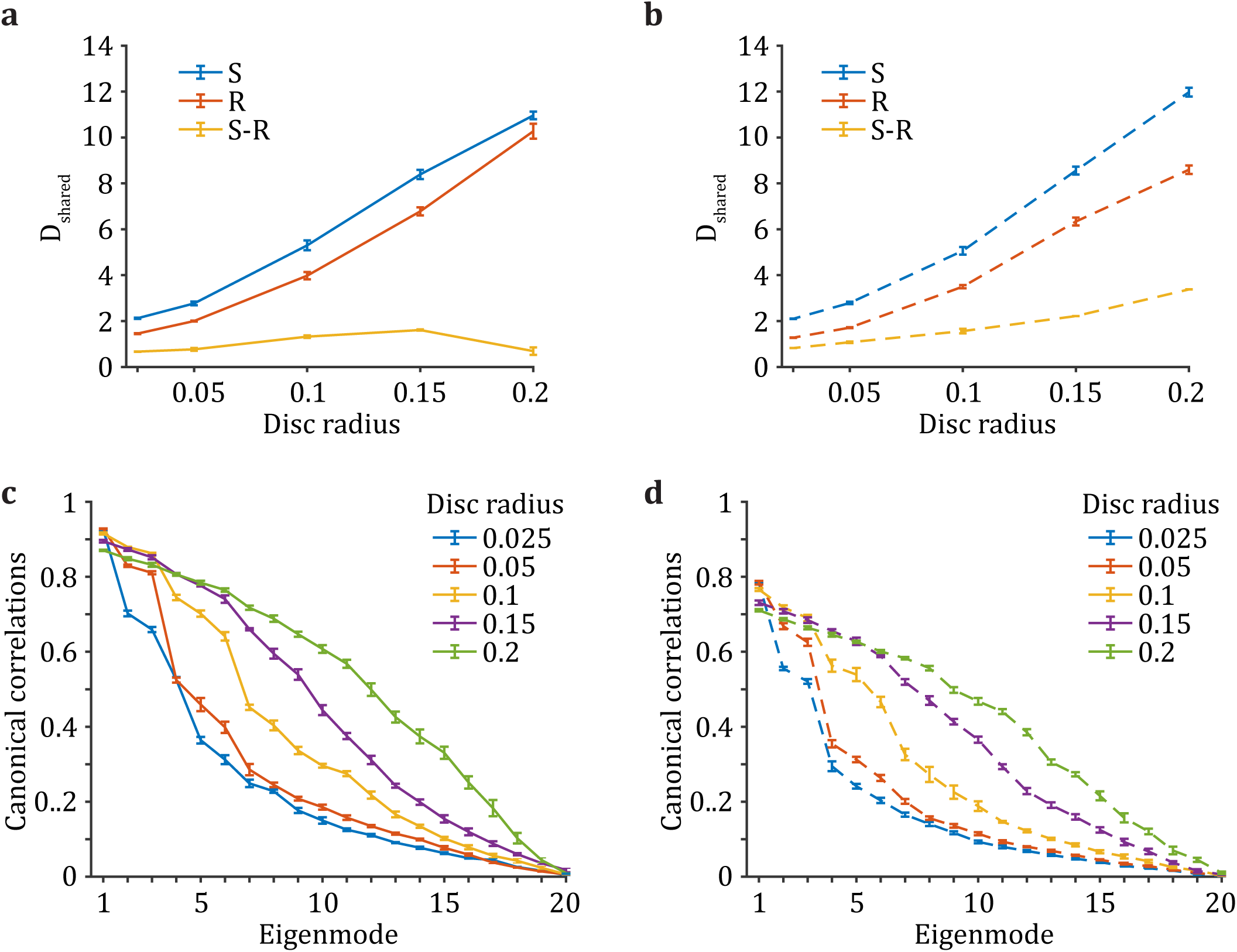
Emergence of novel spatio-temporal patterns in the receiver network induces a misalignment of the shared variability in sending and receiving populations. (**a**) Shared dimensionality *D*_shared_ in the sender network (S), the receiver network (R), and the difference between the two (S-R) in the case where spatio-temporal patterns emerge in S only (Fig. 5a, top). (**b**) Same as (a), except that different spatio-temporal patterns emerge in S and R (Fig. 5a, bottom). (**c,d**) Canonical correlations between the shared fluctuations in S and R when spatio- temporal patterns emerge in S only (c), and when different spatio-temporal patterns emerge in S and R (d).

